# Optimal use of simplified social information in sequential decision-making

**DOI:** 10.1101/2021.01.25.428128

**Authors:** Richard P. Mann

## Abstract

Social animals can improve their decisions by attending to the choices made by others. The rewards gained by attending to this social information must be balanced against the costs of obtaining and processing it. Previous work has investigated the behaviour of rational agents that respond optimally to a full sequence of prior decisions. However, such full sequences are potentially difficult to perceive and costly to process. As such, real animals are likely to rely on simpler forms of information when making decisions, which in turn will affect the social behaviour they exhibit. In this paper I derive the optimal policy for rational agents responding to specific simplified forms of social information. I show how the behaviour of agents attending to the total aggregate number of previous choices differs from those attending to more dynamic information provided by the most recent prior decision, and I propose a hybrid strategy that incorporates both information sources to give a highly accurate approximation to the optimal policy with the full sequence. Finally I analyse the evolutionary stability of each strategy depending on the cost of cognition and perception, showing that a hybrid strategy dominates when this cost is low but non-zero, while attending to the most recent decision is dominant when costs are high. These results show that agents can employ highly effective social decision-making rules without requiring unrealistic cognitive capacities, and point to likely ecological variation in the social information different animals attend to.

## Introduction

Living in groups conveys many advantages for social animals [1, 2]. One of these is the ability to make use of information provided by other individuals in the group. For example: a navigating bird may choose its direction of flight in part by following others in the flock [3, 4]; foraging animals can use the presence of others at a site as an indicator of food availability [5, 6]; fish can effectively avoid predators by following others who are seeking to do the same [7, 8]; and demand for consumer items is often driven by their perceived popularity [9], especially amongst high-status individuals [10].

Substantial research effort has been directed to identifying precisely what types of social information that humans and animals attend to, and the rules that translate that information into behaviours. These studies broadly fall into two (not mutually exclusive) categories: data-driven analysis of the social correlates of behaviour (e.g. [11, 12, 13, 14, 15, 16, 17]) and model-based inference of explicit rules of interaction (e.g. [18, 19, 20, 21, 22, 23, 24]). The first approach is limited by the fact that correlations may result from non-causal relationships between observable social features and behaviour. For example, the orientation of one animal may be correlated with many others in the group beyond those it directly interacts with, since those individuals also interact with others. Furthermore, even the most sophisticated data-driven analysis (e.g. [17]) can only link what is measured, and choosing these measurements requires biological insight. The second methodology proposes causal models to explain observed behaviours, and selects between these by assessing some measure of model fit. To the extent that these models represent a valid causal pathway from social information to behaviour, this avoids the problem of spurious or misleading correlations. However, since the set of all possible causal models linking even one source of social information to behaviour is far too great to test exhaustively, model-based inference is limited in scope unless one can systematically reduce the set of plausible models using *a priori* biological reasoning.

What should that biological reasoning be? I argue that optimality should bridge the gap between information and behaviour. In social animals that are routinely exposed to the choices others make, responses to that information source will be subject to selection [25, 26] and thus are likely to be optimised with respect to fitness. Agents who do not respond optimally to the available information should be out-competed by those who do, whether by evolved instinct or the capability to learn, depending on the life cycle of the species. This implies that individuals’ responses to social information should be understood by considering their adaptive value through rational decision-making theory [27]. However, advanced cognition and accurate perception impose significant costs that create a selective pressure against behaviours that require either, and favouring simplified perceptual models and cognitive processes [28, 29]. Therefore optimal use of social information should be understood as a balance that can vary across species or contexts as these costs change.

Recent theoretical work on sequential decision-making identified the optimal response of an agent to observing a sequence of prior decisions [30, 31]. These studies showed that rational use of information from previous decisions makes use of full sequence information. In particular, in common with earlier models [32, 33], this research shows that recent decisions should be given greater weight by the focal agent when deciding who to follow. However, it is unlikely that real animals have the perceptual acuity, cognitive capacity and time to either deductively apply the complex recursive calculations necessary in these models, or to inductively learn or evolve appropriate responses to every possible sequence they may observe. Instead, it is likely they apply simpler rules based on reduced representations of the sequence information. One such possibility is to respond to aggregate information: the number of agents choosing each option, independent of sequence. This is the basis for many ‘quorum’ models of collective decision-making [34, 35]. Another is indicated by a recent study by Kadak and Miller [24], which confirmed the importance of sequence-dependent decision-making but showed that a ‘follow the last choice’ heuristic was better able to predict decisions in zebrafish than more complex models. These simple rules allow animals to forgo significant cognitive and perceptual costs, but they remain subject to optimisation by selection.

In this paper I identify the optimal rules of social information use by identical agents with a restricted view of previous decisions. I analyse the micro-scale individual decisions and macro-scale collective behaviour of agents rationally responding to different forms of social information and thus assess what information is necessary in order to approximate the behaviour of agents responding to full decision sequences. While varying the cost of cognition and perception I analyse the expected rewards of different social information strategies in competition with each other and thus identify the evolutionary stable strategies under different cost regime.

## Results

In this section I identify the optimal behaviour of identical rational agents attending to restricted subsets of the available social information in a binary choice scenario. Specifically I focus on the use of aggregate information (the number of previous decisions made for each of the two available options) and dynamic information (the choice made by the most recent decision maker). I analyse the optimal behaviour by mapping the observable social responses (the probability that a focal agent chooses a specific option, conditioned on the social context), and how well each strategy fits the behaviour of agents with full, unrestricted social information. Here I derive the consequences of the optimal decision rule both for individual decision makers and at the collective level. In all cases I consider a symmetric choice scenario where the true utility difference between the two choices is zero, and all calculations are performed assuming a fixed environmental noise-to-signal ratio of one (*∊* = 1, see Methods). Previous work has shown that among identical rational agents the value of the environmental noise level does not significantly affect social behaviour [30].

### Aggregate and dynamic social information

The large majority of theoretical and modelling literature on sequential decision-making assumes that agents follow a social feedback rule based on the aggregated number of individuals that have previously chosen each option. The first model of social information I consider is therefore based on this aggregated information. An agent employing this ‘aggregate’ strategy attends to the number of agents that have previously chosen options A or B, and does not observe, or attend to, any information related to the order in which these decisions were made.

Standing against a rule based solely on aggregated previous decisions there is both theoretical and empirical evidence for the greater importance of more recent decisions. In particular, the most recent decision can override a large majority of aggregate decisions in favour of another option [30, 24]. Hence, I also consider how agents should respond when they are only able to observe the most recent decision, and not the aggregate number of previous decisions. Given that this information can rapidly change relative to the aggregate number of decisions, I call this ‘dynamic’ social information, following the same terminology in ref [20].

For each of these two scenarios I calculate how an agent should rationally respond to the social information they are presented with in order to maximise their expected utility, making the assumption that agents are aware that decisions have been made sequentially (although they cannot observe the full sequence), and that other agents are subject to the same limitation as themselves. I focus on identical agents (those sharing the same utility function and cognitive processes) as previous research has shown that agent preferences must be strongly aligned to produce significant social information use [31], and this assumption also permits a more straightforward mathematical treatment. These calculations (see Methods) result in a single number associated with each possible distinct observable social information, which denotes a critical threshold for the agent’s private information: if their private information exceeds this threshold they will choose one option, otherwise they will choose the other.

### Observed social response

Based on these calculations, I analysed all possible outcomes for a group of 10 agents making sequential binary decisions. To analyse these outcomes I first imagine making an empirical study of the observable decisions in the manner of previous attempting direct data-driven inference of social interaction rules (e.g. [12, 13, 15]). To do this I record, for each decision, the aggregate number of previous decisions for options A and B, the identity of the most recent decision and the resulting proportion of times the focal agent chooses each option itself. The result of this analysis is shown in Figure 1, where panels are arranged in two rows: the top row shows the probability for the focal agent to choose option A, conditioned on only the aggregate number of previous choices *n_A_* and *n_B_*, while the lower row shows the same information further conditioned on knowing that the most recent decision maker chose option A. These results are further divided in columns according to each model, with the aggregate model shown in panels A and B, and the dynamic model in panels C, D. For comparison, the equivalent analysis performed on the full sequential model of ref. [30] is shown in panels G and H. These results show a relatively close visual match between the aggregate model and the sequential model when viewed independently of the most recent decision, though the social reinforcement in the aggregate model is somewhat stronger than in the sequential model compressed into this aggregate projection. By contrast, the dynamic model shows a very different pattern: the probability that the focal agent will choose A is approximately independent of *n_A_* and *n_B_* except in the case where one of these values is zero. This result, if seen experimentally, could be interpreted as evidence that the agents are principally concerned not to be alone, and are otherwise content to choose any option that has already been chosen by at least one other agent. However, by construction it is clear this is not the case; instead agents are attending to a different form of social information that is not captured by this aggregate viewpoint. The lower panels show how the apparent social interaction rules change if we include this dynamic social information: the aggregate model remains unchanged (again, by construction), while the dynamic model shows a strong, uniform tendency to follow the most recent decision. The full sequential model also shows a substantial impact of dynamic social information, modulated somewhat by the aggregate values of *n_A_* and *n_B_*. Therefore, not only does the type of social information an agent attends to affect the social interactions it exhibits, but additionally the way that social information is measured determines which model is apparently closest to the behaviour of agents with full sequential information.

**Figure 1:**
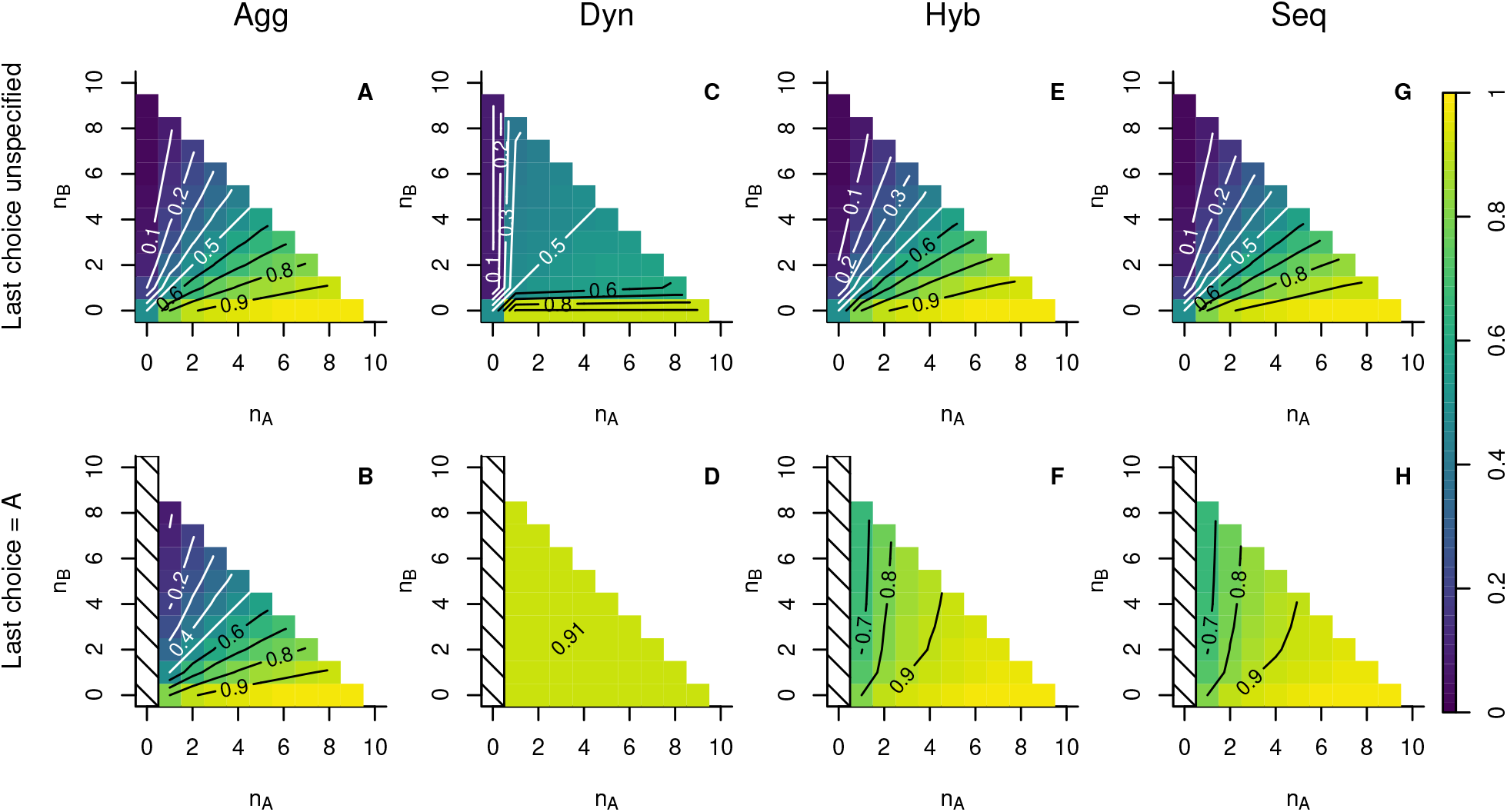
Observed social responses by agents employing different social information strategies. Top row: the probability the focal agent will choose A, conditioned on the number of previous decisions for A and B. Bottom row: the probability that the focal agent will choose A, conditioned on the number of previous decisions for A and B, when the most recent decision was for A. Panels are organised by column for each strategy: (A,B) aggregate strategy; (C,D) dynamic strategy; (E,F) hybrid strategy; (G, H) full sequential strategy.

### Model selection

Secondly I imagine performing a model selection study, assessing how well the aggregate and dynamic models fit the behaviour of agents with access to the full sequential information at the individual and collective levels [36]. First I measure how closely a limited agent approximates the decisions made by one with access to the full sequential information. Assuming that agents actually attend to the full sequence of decisions (using rules as derived in ref [30]), I evaluated the probability assigned to their decisions by both the aggregate model and the dynamic model. This likelihood (the probability of the data, conditioned on a model), shown in Figure 2 A, indicates which model would be favoured by a individual-level model selection analysis (see e.g. [37, 19, 20, 38, 23, 24]). The results show that the dynamic model (blue triangle) is assigned a higher likelihood than the aggregate model (green square). Since an arbitrary number of hypothetical experiments could be considered, I show the likelihood per decision (i.e. the geometric mean probability assigned to each observed decision). I then evaluate the distribution of group outcomes: the total number of agents choosing one of the two options. Comparing this between the limited information models and the full sequential model shows to what degree the two produce similar collective behaviours. Figure 2 B shows that agents attending to aggregate information (green squares) produce a collective outcome that is much closer to that produced from full sequence information (black circles) than that produced by those attending to the most recent decision (blue triangles). While all models show consensus as the most probable outcome, this tendency is weaker in the dynamic model, where intermediate outcomes are all more equally probable than in both the sequential and aggregate models.

**Figure 2:**
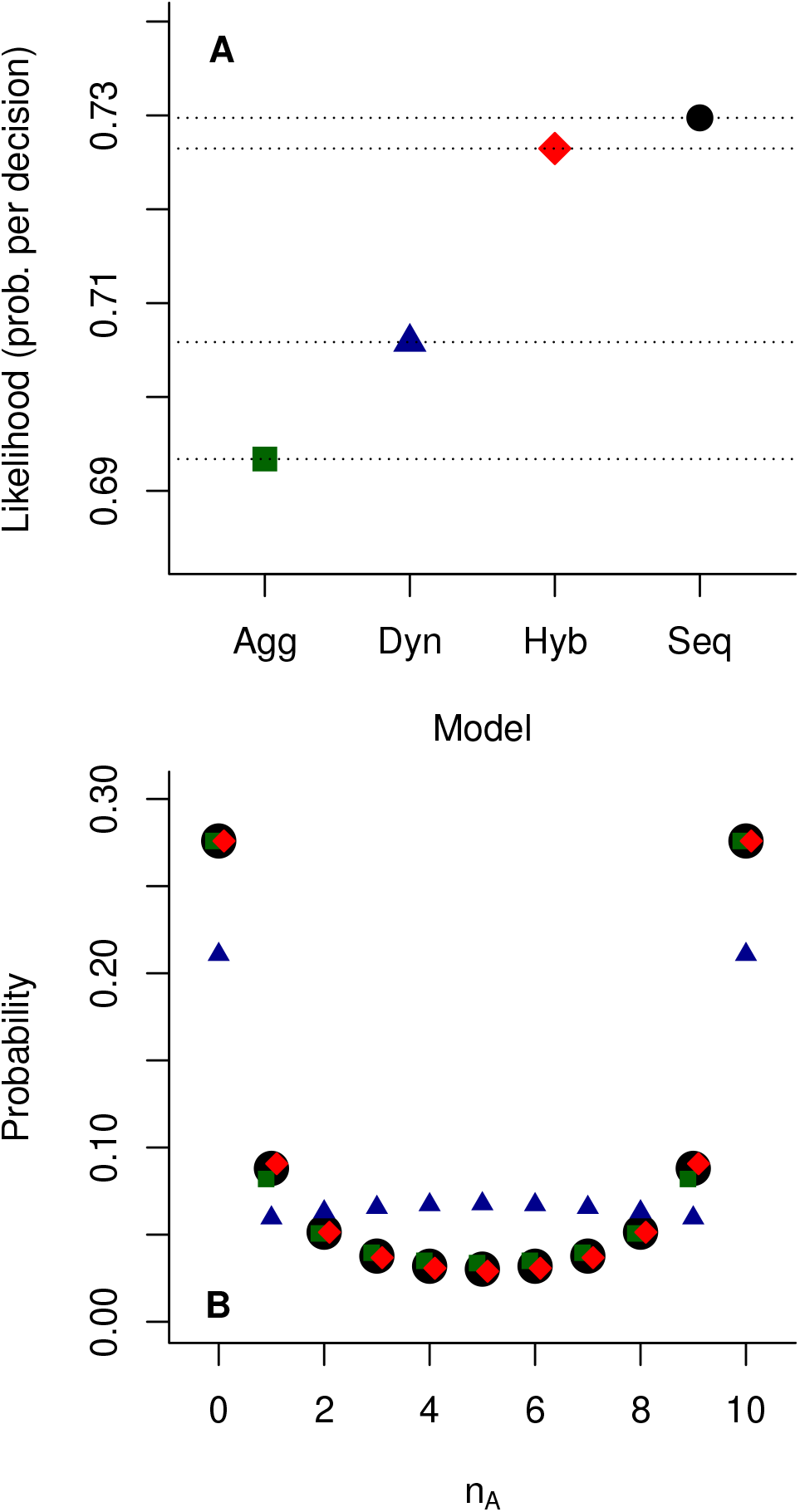
Goodness-of-fit between each limited social information strategy and agents with full sequential information. Panel A shows the likelihood (probability per decision) assigned by models of each strategy to decisions made by fully-informed (sequential strategy) agents. Panel B shows the probability distribution of different collective outcomes (number of agents choosing A) under each strategy.

### Hybrid strategy

The results above pose a problem: the aggregate model produces a good approximation to observed collective behaviour in the sequential model and closely matches the sequential model when social information is measured in terms of aggregate quantities *n_A_* and *n_B_*, while the dynamic model does not. However, the dynamic model more closely predicts the results of individual decisions, and incorporates known facets of recent empirical research, as well as the importance of recent decisions identified in full sequential decision making [30]. When dynamic information is incorporated in the view of the sequential model, the resulting measured social interactions incorporate features of both the aggregate and dynamic models (Figure 1H). It is natural therefore to consider a hybrid model in which agents attend to both the aggregate numbers and to the most recent decision. In a hybrid model, a focal agent may respond differently to the same aggregate configuration of previous decisions depending on the choice made by the most recent decision-maker. This potentially allows the agent to respond to the most important element of the decision sequence, without needing to observe or attend to the full sequence. As shown in Figure 2 A, this hybrid model (red diamond) is assigned a substantially higher individual-based likelihood than either the aggregate or dynamic models, and nearly matches the maximum achievable likelihood as given to the full sequential model (black circle). On the group level, as shown in Figure 2 B, the hybrid model (red squares) achieves an almost perfect match to the collective outcomes from the full sequential model. Having established this superior performance of the hybrid model in matching the individual and collective behaviour of groups of agents with full sequential information, we can also return to the analysis of Figure 1 and ask what the implied observable social interactions of the hybrid model are. Here too we see that the hybrid model is a substantially better match to the sequential model than either the aggregate or dynamical models, both when the most recent decision is ignored (panel E) or incorporated (panel F) in the analysis.

### Evolutionary stability

In deriving the decision rules of agents attending to either full sequential information [30] or the limited social information models presented in this paper, a crucial mathematical assumption is that these decision processes are self-consistent. That is, the decision rule applied by an agent is the rational, optimal action if all other agents employ the same cognitive process and attend to the same information. However, in order for this assumption to hold, the relevant model employed by the agents must either be forced by external factors (if, for example, agents are unable to observe some feature of the social information by virtue of an environmental constraint), or the strategy must be resistant to invasion by agents employing a different cognitive model.

To assess the evolutionary stability of the different social information strategies, I created a scenario where a single individual following one strategy attempts to invade a population composed of individuals following another. In this scenario, I calculate the expected reward gained by the invader and the rest of the population, integrating over the full range of possible true utility differences (*x*) and assuming that the invading individual is equally likely to find itself at any position within the sequence of decisions. Figure 3A shows the difference in reward between the invader and the population for all pairs of strategies, as a percentage of the expected reward obtained by the population in the absence of an invader (note, this matrix need not be symmetric). Positive values (colour coded blue) indicate that the invader gains a greater reward than the population and thus a successful invasion, and negative values (red) indicate an unsuccessful invasion. As expected, the full sequential model, taking account of all information, can successfully invade all other strategies, and is universally resistant to invasion. The hybrid strategy can successfully invade both the dynamic and aggregate strategies. Notably, the dynamic strategy can invade the aggregate strategy, while the aggregate strategy can be successfully invaded by any other. This is despite the aggregate strategy having substantially greater complexity, in terms of the number of distinct social information configurations, than the dynamic strategy. Although the full sequential strategy gains higher rewards than any competing strategy, this comes at the cost of high complexity. An agent wishing to employ this strategy must either complete a complex recursive mathematical analysis [30], or otherwise learn how to respond to a large number of possible sequentially-ordered social information configurations; assuming that that no symmetry principles are employed to reduce the number of distinct possibilities there are 2^*n*^ − 1 possible configurations they might observe. Other strategies also vary widely in complexity, with 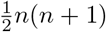 configurations for the aggregate strategy (each possible combination of *n_A_* and *n_B_*), *n*(*n* − 1) + 1 for the hybrid strategy (the same configurations with either two possible choices for the most recent decision, excluding impossible configurations) and just 3 for the dynamic strategy (two possible most recent decisions, or no social information). Algorithmic complexity imposes cognitive costs on agents, whether through the energy cost of maintaining sufficient neurological and sensory capacity to employ the algorithm or in the additional time it takes to learn a complex response. Therefore I reevaluated the above evolutionary stability analysis for a range of hypothetical cognitive costs, defined as a fixed cost per distinct configuration an agent must learn to employ the strategy. These hypothetical costs are chosen to illustrate three distinct regimes: (i) for a low but non-zero cognitive cost, the hybrid model is dominant (Figure 3B), successfully invading all other strategies and resisting invasion by all; (ii) with an intermediate cognitive cost, hybrid and dynamic strategies can coexist – each can invade the other when the other is dominant, and resist invasion by the sequential or aggregate strategies (Figure 3C); (iii) for sufficiently high cognitive costs the dynamic strategy is dominant (Figure 3D), as a result of its very low algorithmic complexity. There is no value for the cognitive cost that will make the aggregate strategy evolutionarily stable, since it always out competed by the dynamic strategy.

**Figure 3:**
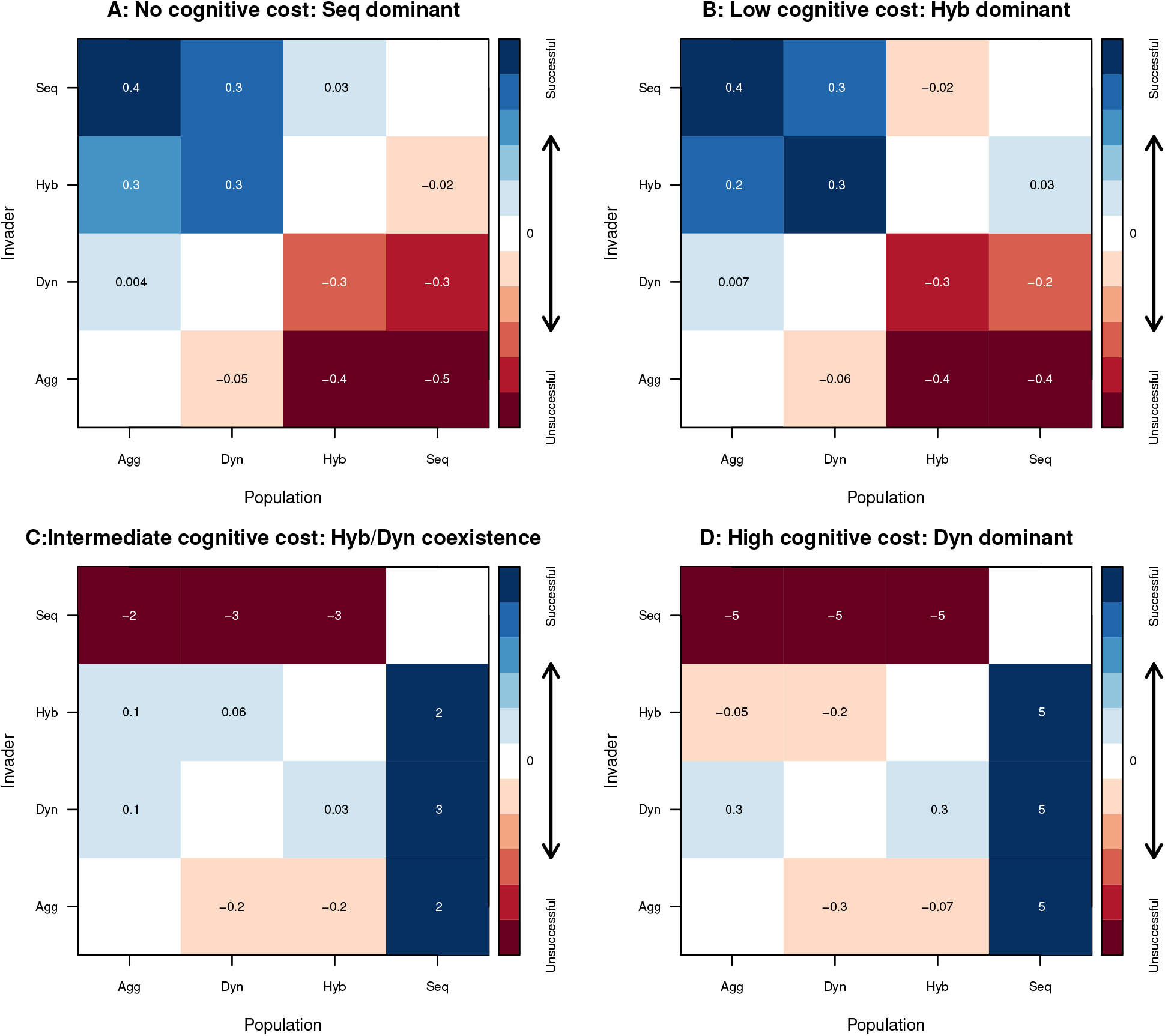
Evolutionary stability analysis. Panel A shows the percentage difference in expected rewards for a single individual of each strategy invading a population of agents employing a different strategy, in the absence of any cognitive costs. Positive quantities (blue) show that the invader gains greater rewards than the other agents. Panels B-D show a progression of increasing cognitive costs (defined as a fixed cost per distinct possible configuration of social information), showing a different single or mixed evolutionarily stable strategy for differing costs. These costs are given as a proportion of the typical reward scale (the standard deviation of *x*), and are for each case: (A) 0; (B) 4 × 10^−7^; (C) 2 × 10^−5^; and (D) 4 × 10^−5^.

## Discussion

I derived the optimal policy for agents attending to limited social information in a sequential decision-making setting, and analysed the resulting behaviour of the agents at the individual and collective level. All three decision rules based on limited social information are substantially simpler than the previously developed full sequential decision rule, when viewed in terms of the number of distinct configurations of social information that an agent must be in a position to respond to. This implies that while deductive calculation of the rational behaviour remains complex, inductive learning or adaptation is more straightforward than when attending to full sequential information. Although the application of symmetry principles and parametric function approximations may change the quantitative complexity of each strategy, the ordering of complexity is likely fixed, with dynamic social information (the most recent decision) being the simplest and fully ordered sequences of previous decisions the most complex.

The results presented in this paper reveal two key insights. First, the behaviour of agents attending to full sequential social information can be approximated to a very high degree by a simpler strategy that makes use of only the aggregate number of previous decisions for each option, along with the identity of only the most recent decision. That is, almost all the information contained in the specific ordering of decisions, above the aggregate number of choices, is contained within the most recent decision. Employing this ‘hybrid’ strategy allows agents to make nearly identical decisions to those with full sequential information without employing complex mathematical calculations, and to obtain greater rewards than two simpler strategies making use of only the aggregate quantities or the most recent decision alone. Second, attending to different forms of social information produced differing expected rewards for agents, and the ordering of these rewards did not correspond to the relative complexities of the forms of social information. In particular, attending to just the most recent decision produced greater rewards than attending to aggregate information when the two strategies were in competition.

While specific environmental or cognitive limitations may force individuals to attend to a particular feature of social information, in many cases agents can be expected to evolve the capacity to attend to information that increases their expected reward relative to competitors. In such cases the hybrid model, wherein an agent attends to both the aggregate numbers of previous decisions across different options and the identity of the most recent decision (but no other sequence information), provides a highly accurate approximation to the full sequential model, with a dramatically lower cognitive cost. It also provides a clear improvement over either aggregate or dynamic social information alone. Given the very high complexity of the full sequential model, we should therefore expect such a hybrid model to be favoured when the cost of cognition or perception is relatively small but non-negligible. When cognitive or perceptual costs are very high, it is likely that a dynamic social information model would be favoured if this information is available – this model has very little cognitive complexity and outperforms the aggregate model for any cognitive cost. The results presented here also suggest that there may be scenarios in which no single strategy is dominant. In this case agents may specialise into multiple co-existing social information strategies. In the example shown here (Figure 3C), agents employing a hybrid strategy would obtain a somewhat higher reward than agents employing a dynamic strategy, that balanced the higher cognitive cost they incur. This is analogous to the emergence of ‘informed’ and ‘uninformed’ agents in collective navigation [4, 39]. Because the strategies explored in this paper are only optimal assuming that every agent employs the same strategy, further research is needed to determine how these strategies would adapt under co-evolution.

Aggregate social information, which forms the basis of many models studied in the literature (e.g. [40, 41, 34, 35, 42, 43, 44]), is outperformed by at least one other model for any cognitive cost (under the assumption that cognitive cost varies monotonically with the number of distinct social information configurations for which a threshold must be learned). Although the quantitative complexity of this strategy could be reduced by exploiting symmetries and by approximating the social response with a parametric representation, it is unlikely that this could reduce the complexity below that of the dynamic model. It is likely that in some contexts individuals may be limited to this information by environmental or other external constraints, such as ants attending to deposited pheromones as a proxy for the number of individuals making each choice [40, 41, 14] or audience members attending to the volume of applause as a proxy for the number of individuals clapping [21]. However the poor performance of aggregate social information should serve as a check on the future use of such models without considering alternatives. Where an aggregate social information model is found to be the best fit to some empirical data, this must either be the result of some external constraint on the observable social information or a violation of one or more assumptions in this paper. For instance, the models presented here treat the choices of others as valuable knowledge solely through the information they transmit about the utilities of the possible options. An intrinsic benefit of aggregation, such as predation-dilution effects [45, 1], could make knowledge of aggregate quantities useful beyond the direct social information value.

The results of this study have both a descriptive and a normative interpretation. They indicate how the use of social information is likely to have evolved when faced with cognitive limits on the complexity of social information that can be considered. They also suggest how we should respond to the decisions we observe others making: by attending to both aggregate and dynamic social information. We should be wary of options that have been chosen by many other individuals, but which few individuals have chosen recently; current unpopularity may indicate that new information has become available that outweighs the aggregate social information in favour of the choice. It is reasonable to expect that the most recent decisions may often be available to us when making a decision, even when the full sequence is not observed, and the results here suggest that combining these recent decisions with aggregate quantities is sufficient to gain most of the possible advantages of social information.

In this study I have modelled both environmental and social information as abstractions, without considering how perception and cognition are achieved on a neurological level. Whereas I have modelled the cost of perception and cognition in simple terms, further research on the neural correlates of social decision making may reveal specific forms of social information that are inherently more favourable for neurological representation. I have considered only strict subsets of the available social information as possible perceptions. A fascinating possibility is that individuals may evolve perceptual models that do not correspond in any direct way to physical reality, but which are instead adapted to act as ‘user interfaces’ with the world for the purpose of maximising fitness [28, 46]. Future research might investigate this possibility with a view to understanding how social information is perceived on a conscious level, in both humans and animals.

## Methods

I consider a group of identical agents faced with a sequential, binary decision, each choosing between option A and option B. These options have true utilities *U_A_* and *U_B_*. I denote as *x* the true difference in utilities between these options: *x* = *U_A_* − *U_B_*. Each agent seeks to choose the option that maximises its utility, based on its beliefs about *x*. That is:

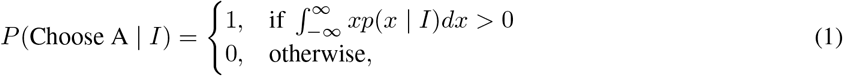

where *I* represents all the information, both private and social, that an agent has.

Each agent begins with an identical prior distribution over the values of *x*; I assume this to be a normal distribution centred on zero (no prior bias for either option) and without loss of generality scale all utilities such that the variance of this distribution is one:

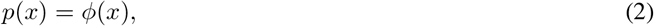

where *ϕ*(·) is the standard normal probability density function.

### Private information

Each agent *i* receives private information Δ_*i*_ about the value of *x*, drawn from a normal distribution with variance *∊*^2^:

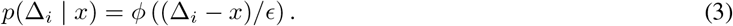

Using their private information, each agent can independently update their belief about the value of *x*, using Bayes
rule:

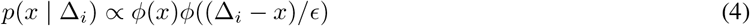

### Social information

Each agent can further update their belief about the value of *x* by conditioning on the available social information available to them, *S_i_*. The general form of this update again utilises Bayes rule:

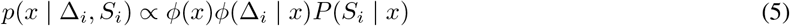

The structure of *S_i_* and the resulting form of *P* (*S_i_* | *x*) differs when considering the different types of social information that agent may attend to. For the case where an agent attends to the full sequence of previous decisions, *S_i_* = {*C*_1_*, C*_2_, … *C_i_*_−1_}, where *C_j_* is the choice made by agent j (*C_j_* = 1 or −1 if agent j chooses A and B respectively). In this case, ref [30] derives a recursive expression for *p*(*x* | *C*_1_, …, *C_i_*_−1_, Δ_*i*_)

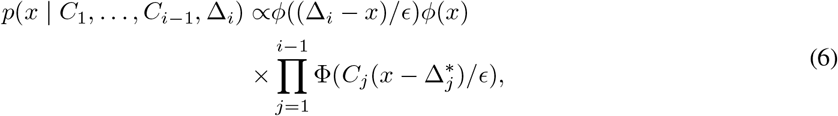

and

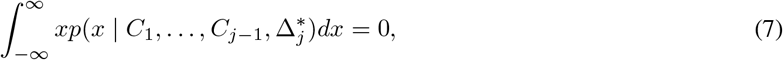

where 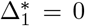. The values of 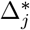 represent a series of critical thresholds; each agent *j* chooses A if and only if 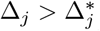. Since these thresholds can in principle be calculated by any other agent, this allows later agents to identify whether the private information of agent *j* was above or below this threshold. Hence, every possible sequence of previous decisions is ultimately associated with a critical threshold, which in turn determines that probability that the agent will be observed to select option A or B through the stochastic generation of Δ_*j*_ from the environment.

### Observation

In common with previous developments of this model [30, 31], I evaluate the probability of an agent choosing A or B by considering the position of an external observer, for whom the private information of all agents is unknown. In this paper I assume that all observations are made under natural conditions, meaning that the experimental noise level is the same as the habitual noise level (see ref [30] for a discussion on the role of experimental noise). The external observer can calculate the critical information threshold, Δ* for any agent by the process laid out above. Having evaluated this threshold, the probability that the agent will choose option A is the probability that the agent’s private information exceeds this threshold.

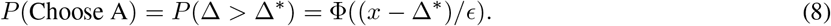

### Simplified social information

When agents attend to simpler social information this represents a compression of the full sequence of decisions into a lower-dimensional projection, and hence this is a many-to-one relationship; multiple full sequences may be compatible with the observed social information. For example, if an agent sees that one agent has already selected A and one has selected B, this is compatible with the first decision being for A and the second for B, or vice versa.

Ultimately an agent must associate each distinct observable social information with a threshold, as with the distinct observable sequences above. This threshold specifies that value of private information Δ_*i*_ that would make the expected value of *x* zero. These thresholds could in principle be acquired through evolutionary adaptation or reinforcement learning, but in order to calculate them from first principles one must imagine the agent performing a complex cognitive process that depends on the precise nature of the simplified social information, as detailed in the specific cases below.

### Aggregate and hybrid strategies

. I take as the first case an agent that observes the aggregated number of decisions for A and B (*n_A_* and *n_B_* respectively), but without any information regarding the order of those decisions. Consider the set 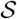 of all sequences of decisions that could have led to this configuration (of which there are (*n_A_* + *n_B_*)!/*n_A_*!*n_B_*! possibilities). Each sequence 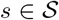 specifies a series of previous choices *C_k_* ∈ {1, −1}, *k* = 1, …, *n_A_* + *n_B_*, and associated aggregate configurations (previous values of *n_A_* and *n_B_*). Denoting the associated thresholds for these configurations as 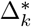 within each sequence *s*, then agent *i*’s belief over *x* is a weighted mixture of the appropriate beliefs for each possible sequence:

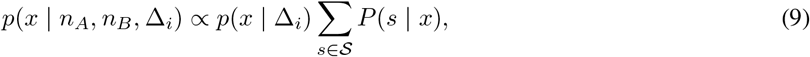

where:

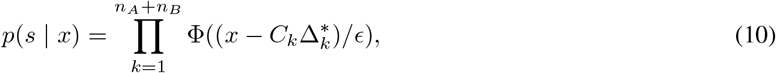

is the probability of the specific sequence *s*, conditioned on the true utility difference *x*. Note that the threshold values within each sequence, 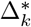, are not the same values as for that sequence within the full sequential model (i.e. in equation 6); instead they are thresholds specific to the aggregate configurations *n_A_*, *n_B_* that will have resulted from the (*k* − 1)th choice along sequence *s*.

Similarly to equation 7, this then recursively specifies the appropriate threshold 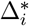 for the configuration *n_A_, n_B_*:

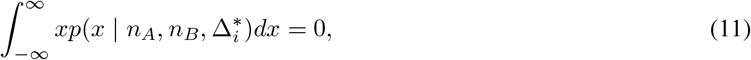

An analogous process can be followed for the hybrid model, by considering all sequences that agree with the most-recent decision and the aggregate values of *n_A_* and *n_B_*.

### Dynamic strategy

In the dynamic model, an agent attends only to the most recent decision. Therefore the set of possible distinct configurations of social information is limited to three possible observations: a decision for A, a decision for B, or no decision (if the focal agent is the first decision maker). This creates a specific problem for deriving the appropriate critical thresholds: the social information that the focal agent observes may be the same as the previous agent to which it attends. This makes the problem self-referential in a way that the aggregate and hybrid models are not.

Consider the belief of a focal agent *i* about the value of *x*, having observed the previous decision *C*_*i* − 1_ = 1 (i.e. a choice of A) and having private information Δ_*i*_:

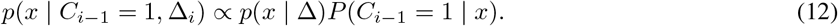

Similarly to equation 9, this can be expressed in terms of the probability *P* (*s* | *x*) of all sequences 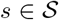 that satisfy *C_i_*_−1_ = 1.

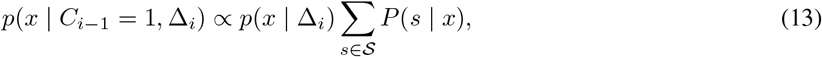

Note that these may be of differing lengths of up to *n* − 1 prior choices; the focal agent can only be aware that they are not the first decision-maker. Assuming that some threshold Δ* exists such that:

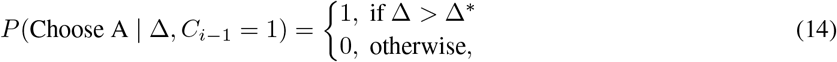

then the probability of a given sequence *s* = {*c*_1_, *C*_2_, … *C_m_*} is given by:

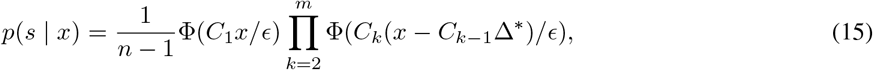

where the initial factor of 1/(*n* − 1) originates from the agent’s uncertainty over the number of prior decisions, which is equally likely to be anywhere from 1 to *n* − 1. Finally, in order to make the decision rule self-consistent requires that the expectation of *x* be zero when Δ_*i*_ = Δ*, thus:

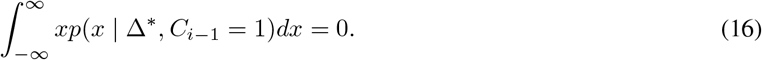

By symmetry, the equivalent threshold for the agent to apply when *C_i_*_−1_ = −1 is −Δ*.

## Acknowledgements

This work was supported by UK Research and Innovation Future Leaders Fellowship MR/S032525/1

